# TBP facilitates RNA Polymerase I transcription following mitosis

**DOI:** 10.1101/2023.07.13.548763

**Authors:** James Z.J. Kwan, Thomas F. Nguyen, Sheila S. Teves

## Abstract

The TATA-box binding protein (TBP) is the sole transcription factor common in the initiation complexes of the three major eukaryotic RNA Polymerases (Pol I, II, and III). Decades of research have shown that TBP is essential for proper transcription by the three RNA Pols, though the emergence of TBP paralogs throughout evolution have expanded the complexity in RNA Pol initiation. We have previously shown that acute TBP depletion in mouse embryonic stem cells led to a global decrease in Pol III activity, consistent with the requirement of TBP in Pol III inititation. In contrast, Pol II transcription remained unaffected in the absence of TBP and its paralogs. In this report, we show that, in contrast to Pol II-transcribed genes, the TBP paralog TRF2 does not bind to Pol I promoters, and therefore cannot functionally replace TBP upon depletion. Importantly, acute TBP depletion has no major effect on Pol I occupancy or activity on ribosomal RNA genes, but TBP binding in mitosis leads to efficient Pol I reactivation following cell division. These findings provide a more nuanced role for TBP in Pol I transcription in murine embryonic stem cells.

**Significance:** The TATA-box binding protein (TBP) is a highly conserved and essential protein in eukaryotes. Decades of *in vitro* and yeast research have established its role in the initiation of the three main eukaryotic RNA polymerases. However, the ability to rapidly deplete proteins *in vivo* is revealing more nuance in the function of TBP in mammalian cells. Using this technology, we reassess the role of TBP in RNA Polymerase I (Pol I) transcription in mouse embryonic stem cells. We find that neither TBP nor its paralog TRF2 are required for Pol I recruitment or activity, but TBP binding during mitosis promotes efficient reactivation after cell division. Overall, these findings provide new evidence into the complex function of TBP in eukaryotic transcription.

## Introduction

Eukaryotic RNA transcription is mainly directed by three distinct complexes: RNA Polymerase I, II, and III (Pol I, II, and III). Pol I is dedicated to transcribing the three large ribosomal RNAs (rRNAs), whereas Pol II largely directs the transcription of protein coding mRNAs, and Pol III is responsible for transcribing transfer RNAs (tRNAs) and small rRNAs (1). Although initiation by each RNA Pol requires different sets of general transcription factors, the TATA-box binding protein (TBP) is uniquely common to all three initiation complexes (2, 3). In general, TBP binds onto DNA promoters to recruit Pol-specific general transcription factors and ensure proper polymerase loading. However, the role of TBP in each initiation complex has become more nuanced over evolutionary time. For example, inactivation or nuclear depletion of TBP in budding yeast has been shown to decrease the transcription activity of all three RNA polymerases (4–7). In mammals, like mouse embryonic stem cells (mESCs) and human HAP1 cells, acute depletion of TBP resulted in no changes in Pol II occupancy and transcription, while Pol III-mediated occupancy and transcription of tRNAs was severely impacted (8, 9). TBP has also been shown to facilitate reactivation of Pol II transcription following mitosis in mESCs (10), though such mitotic bookmarking function remains unclear for Pol I or III transcription. Finally, TBP paralogs have further increased the complexity of TBP-mediated RNA Pol initiation throughout evolution (11–15). In mESCs, the TBP paralog TRF2 binds on to Pol II genes, but does not functionally replace TBP for Pol II-mediated transcription (8), though its role in Pol I or III transcription remains unclear.

Pol I-transcribed rRNA comprises the largest component of RNA, making up to 80-90% of the RNA molecules in eukaryotic cells (16), and are central components of the ribosome machinery that carry out protein synthesis. Decades of research have shown that Pol I initiation begins with the binding of the upstream binding factor (UBF) onto the promoters of rRNA genes (rDNA), organized as hundreds of tandem repeats in multiple chromosomes (17, 18). DNA-bound UBF then recruits the selective factor 1 (SL-1, also known as transcription initiation factor IB, TIF-IB), which is a complex composed of TBP, TAF12, TAF1A, TAF1B, TAF1C, and TAF1D (19–21). Binding of SL-1 then leads to the recruitment of Pol I in complex with Rrn3 to the rDNA promoter to begin transcription (22, 23). *In vitro* experiments with yeast extracts have shown that basal levels of Pol I transcription can be achieved in the absence of TBP, but full activation of rRNA transcription still requires TBP (7). However, overexpression of TBP in yeast cells can overcome depletion of upstream transcription factors to support rRNA transcription, highlighting a nuanced role for TBP in Pol I-mediated transcription *in vivo* (24).

In this study, we revisit the role of TBP in the transcription initiation of rRNAs in mESCs. We report that the TBP paralog TRF2 does not bind onto rDNA promoters and therefore does not functionally replace TBP for Pol I transcription. Furthermore, depletion of TBP in mESCs results in no major effects in nascent rDNA transcription despite a minor decrease in Pol I occupancy. Surprisingly, TBP depletion affects the efficacy of Pol I reactivation following mitosis. Taken together, our data suggests that TBP promotes efficient Pol I reactivation from a mitotic state, but is not strictly required for reaching full Pol I transcription.

## Results

### TBP, but not TRF2, binds onto rDNA genes

The mouse genome contains two TBP paralogs, the *Tbpl1* and *Tbpl2* genes that encode for TRF2 and TRF3, respectively (25–27). In mESCs, only TRF2 is expressed (8). To determine if TRF2 has TBP-redundant or -unique roles in Pol I transcription, we utilized our previously established homozygous mAID-TBP cell line (C64) (8, 10), which allows for rapid depletion of TBP upon indole-3-acetic acid (IAA) treatment (28). In this previous study, TBP depletion was confirmed by performing Cleavage Under Targets and Tagmentation (CUT&Tag), a high resolution chromatin profiling technique (29). Standard analysis using the commonly used mm10 genome enabled quantification of TBP binding and its depletion on Pol II and III genes, but the highly repetitive nature of rDNA rendered such analysis difficult for Pol I genes. The eukaryotic rDNA consists of a repeated array of tandem repeats. Each transcription unit is separated by the Intergenic Spacer (IGS) and is composed of Promoters, Enhancer repeats, External and Internal Transcribed Spacers (ETS and ITS respectively), ribosomal RNAs (18S, 5.8S and 28S), and the Transcription Termination Factor binding sites (TTF1 sites) (30, 31). Recent advances have enabled the construction of the human and mouse genome customized for rDNA mapping (30). We therefore realigned the TBP CUT&Tag data to the recently established mm10-rDNA genome (30) and normalized the signal by RPKM. We observe strong binding of TBP under control conditions at the rDNA spacer and 47S promoter with high reproducibility (Figure 1A, Figure S1A). Additionally, treatment of C64 cells with 6 hours of IAA show a strong depletion of bound TBP on the rDNA promoter, confirming previous findings of TBP depletion for Pol II promoters and demonstrating proper alignment to the custom genome (Figure 1A-B).

**Figure 1.**
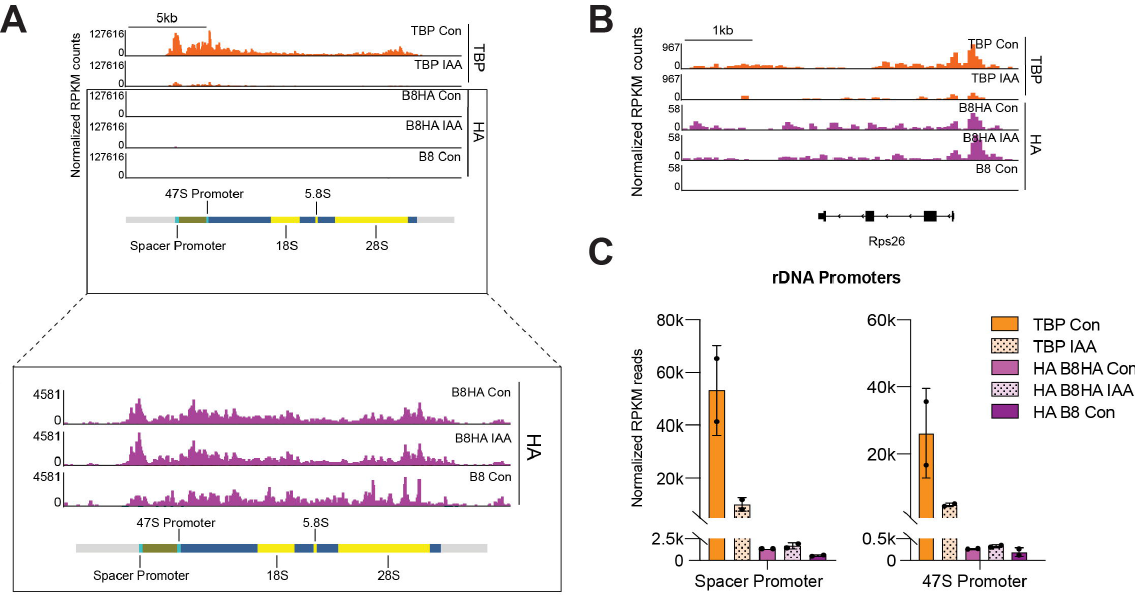
TBP, but not TRF2 binds onto the Spacer and 47S Promoter of rDNA. **(A-B)** Gene browser tracks of the custom rDNA genome on (A) the entire rDNA chromosome (chrR) and (B) the Pol II specific gene (Rps26) for TBP (orange) and TRF2-HA (magenta) in control and indole-3-acetic acid (IAA)-treated C64 mouse embryonic stem cells (mESCs). (**C)** Averaged RPKM normalized read counts binned by 10bp at rDNA Spacer promoter and 47S promoter for TBP control (orange), TBP IAA (spotted light orange), HA B8HA control (magenta), HA B8HA IAA (spotted light magenta) and the negative HA B8 control (violet). Error bars represent standard deviation of 2 biological replicates.

With a system to deplete TBP on Pol I promoters, we next examined the TBP paralog TRF2, first by measuring its binding when TBP is present. We reanalyzed previously published TRF2-HA CUT&Tag data (8). TRF2 knockout mESCs (B8) were used to generate stably overexpressing HA-tagged TRF2 (B8HA), and CUT&Tag was performed on both cell lines using the HA antibody. After aligning to the rDNA genome and normalizing by RPKM, we observe that the TRF2 signal strength on rDNA promoters matches the background noise of the B8 data, suggesting that these mapped reads largely correspond to noise (Figure 1A, Figure S1B). This lack of binding contrasts strikingly with the previously shown ability of TRF2 to bind onto Pol II gene promoters (8), such as the *Rps26* promoter (Figure 1B). We next examined whether TRF2 binding is induced upon TBP depletion by reanalyzing TRF2-HA CUT&Tag data after IAA treatment. We found that depletion of TBP in the B8HA cell line does not increase the binding occupancy of TRF2 at Pol I promoters (Figure 1A-B). To further quantify the CUT&Tag data, we plotted the average TBP and TRF2 signal in the rDNA promoter and gene regions separately, measured by averaging the RPKM normalized reads within 10 bp bins for each region. We observe that TBP occupancy is 10-fold higher at the promoters compared to genic regions, and that IAA-treatment reduced TBP occupancy by ∼80% (Figure 1C, Figure S1C). The levels for HA-tagged TRF2 and the negative control B8 sample remain less than ∼7% of TBP levels throughout all regions (Figure 1C, Figure S1C). Taken together, these results suggest that our previously established mAID-TBP system enables acute TBP depletion in the rDNA genome, and that unlike in mRNA promoters, rDNA promoters are exclusively bound by TBP and not TRF2.

### Effects of TBP depletion on Pol I occupancy and activity

We next examined the effects of TBP depletion on recruitment of Pol I to the rDNA. First, we determined whether TBP depletion affects Pol I protein levels. We performed Western blot on control and IAA-treated C64 whole cell lysates and blotted for TBP, the Pol I subunit RPA1, and the loading control Tubulin. We detect near complete depletion of TBP in the IAA treated samples, but RPA1 levels remain unchanged (Figure 2A), suggesting that acute TBP depletion does not affect Pol I protein levels.

**Figure 2.**
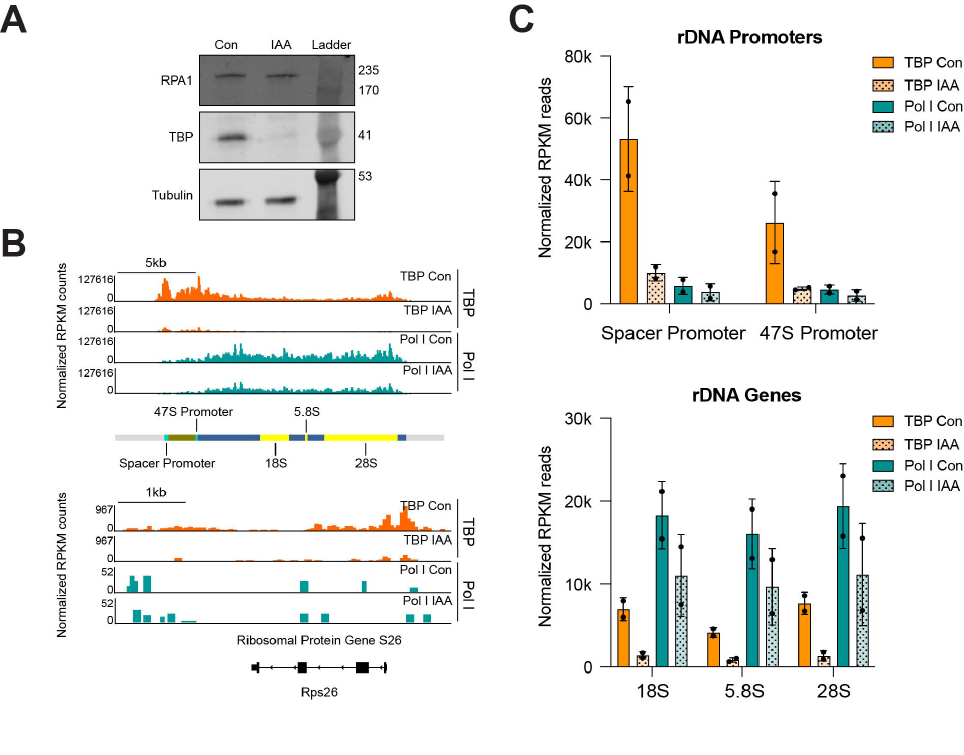
TBP depletion slightly impairs the binding of Pol I on rDNA genes. **(A)** Western blot analyses of whole cell lysates for control and IAA-treated C64 mESCs blotting for ɑ-RPA1, ɑ-TBP, with ɑ-Tubulin used as a loading control. Cells were incubated with DMSO (control) or 500 μM IAA for 6 hours. **(B)** Gene browser tracks of the custom mm10-rDNA genome (top) and the Pol II specific gene Rps26 (bottom) for TBP (orange) and Pol I (teal) in control and IAA-treated C64 mESCs. **(C)** Averaged RPKM normalized read counts binned by 10bp at rDNA promoters (Spacer and 47S) and gene bodies (18S, 5.8S and 18S) for TBP control (orange), TBP IAA (spotted light orange), Pol I control (teal), Pol I IAA (spotted light teal). Error bars represent standard deviation of 2 biological replicates.

To investigate the effects of TBP on Pol I occupancy, we performed Pol I CUT&Tag on control and IAA-treated C64 cells, aligned the data to the custom mm10-rDNA genome and normalized the signal by RPKM. We observed that, in control samples, Pol I binds to the rDNA loci with high reproducibility across replicates, but not at Pol II coding genes such as Rps26 (Figure 2B, Figure S1A), suggesting strong specificity of the assay. To quantify the Pol I signal, we plotted the averaged RPKM read counts mapped at the annotated rDNA promoters and gene bodies. In the promoter region, we detect high levels of TBP and low Pol I occupancy, and within the rDNA gene bodies, we detect low TBP binding and high Pol I occupancy (Figure 2C). Upon TBP depletion with 6 hours of IAA treatment, we observe a 30-40% decrease in Pol I levels on rDNA genes (Figure 2B-C). We then visualized the Pol I occupancy from 5’ start to the 3’ end of each of the annotated rDNA regions as a heatmap binned by 10bp with and without IAA-treatment. We observed high Pol I binding within transcribed regions and no signal at the IGS and TTF1 sites. Furthermore, a consistent 30-40% decrease in Pol I intensity across all Pol I regions in the TBP-depleted sample is detected (Figure S2A), despite no changes in RPA1 protein levels. This consistent decrease, however, is not proportional to the 80-90% decrease in TBP binding, suggesting that our data confirm previous *in vitro* studies showing that TBP is not strictly required for Pol I recruitment and binding to rDNA genes.

CUT&Tag analysis provided a measure of Pol I occupancy on rDNA. To directly measure Pol I transcriptional activity on rDNAs, we analyzed previously published native elongating transcript sequencing (NET-seq) datasets for control and TBP-depleted cells by aligning them to the custom mm10-rDNA genome (8). NET-seq captures nascent RNA from elongating RNA polymerases at single nucleotide resolution and can also capture the activity of elongating Pol I (34). We also re-analyzed published data from chromatin-associated RNA-seq (chrRNA-seq), an orthogonal approach that captures newly transcribed chromatin-associated RNA through biochemical fractionation, performed on C64 mESCs in control and TBP-depleted conditions (10). We normalized the NET-seq and chrRNA-seq signal to spike-in controls and detected comparable signals for control and TBP-depleted cells in the internal transcribed regions, 18S, 5.8S, and 28S rRNAs (Figure 3A). To further quantify the signal, we averaged the normalized NET-seq and chrRNA-seq reads across the intronic and coding rDNA regions, plotted the logged values as a bar plot, and did not observe any significant differences in rRNA read counts for control and TBP-depleted cells (Figure 3B, Figure S3B). We also quantified NET-seq and chrRNA-seq read counts for annotated Pol I rRNAs obtained from the UCSC genome browser (https://genome.ucsc.edu/) (35), displayed the logged counts as a scatterplot, and observed virtually no changes in rRNA reads upon TBP depletion (Figure 3C). Therefore, despite the 30-40% decrease in Pol I occupancy as observed by CUT&Tag, NET-seq, and chrRNA-seq analyses indicate that TBP is not required for proper Pol I transcription of rRNAs in mESCs, in contrast to previous *in vitro* studies.

**Figure 3.**
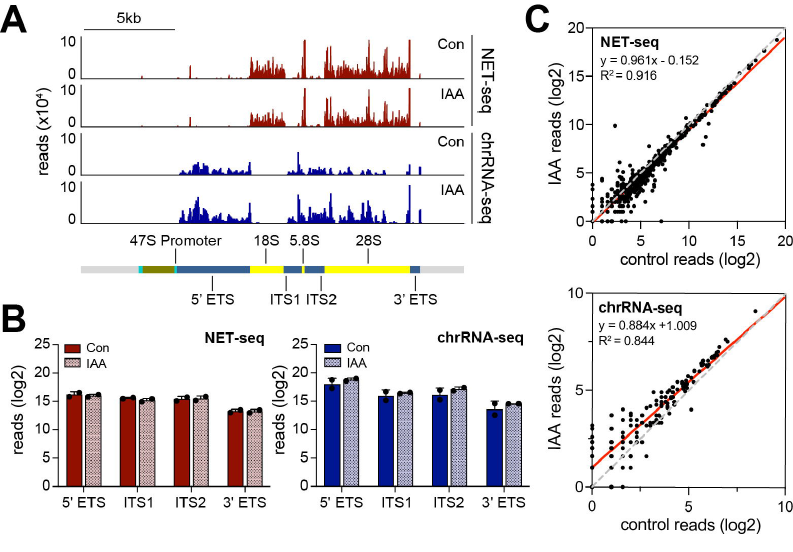
TBP depletion does not affect nascent transcription of rRNAs. **(A)** Gene browser tracks of the custom mm10-rDNA genome chrR for NET-seq (maroon) and chrRNA-seq (blue) in control and IAA-treated C64 mESCs. **(B)** Normalized intronic read counts of NET-seq (left) and chrRNA-seq (right) for control and IAA-treated mESCs across the 5’ external transcribed spacer (5’ ETS), internal transcribed spacers (ITS1 and ITS2), and 3’ external transcribed spacer (3’ ETS) regions. Error bars represent standard deviation of 2 biological replicates (**C**) Normalized read counts of NET-seq (top) and chrRNA-seq (bottom) across annotated Pol I rRNAs for control vs. IAA-treated C64 mESCs.

### Mitotic binding of TBP on rDNA promotes efficient transcriptional reactivation

Similar to Pol II, Pol I transcription is globally inhibited during mitosis (40, 41). Given that TBP has been shown to maintain stable binding at Pol II promoters during mitosis, which is required for efficient reactivation of Pol II genes, we sought to examine the role of TBP in Pol I transcription following mitosis. We first reanalyzed previously published TBP ChIP-seq data in C64 asynchronous and mitotic cells. After normalizing by RPKM and realignment to the rDNA genome, we observe a much stronger binding of TBP to the rDNA genome in mitotic versus asynchronous cells (Figure 4A), with binding levels increased by 10-fold. In contrast, the global average of TBP binding for Pol II genes has been shown to be slightly decreased during mitosis compared to asynchronous cells (10), consistent with our re-analysis on *Gapdh* and *Myc* (Figure S4A). These results suggest that TBP may bookmark rDNA genes more strongly than Pol II genes.

**Figure 4.**
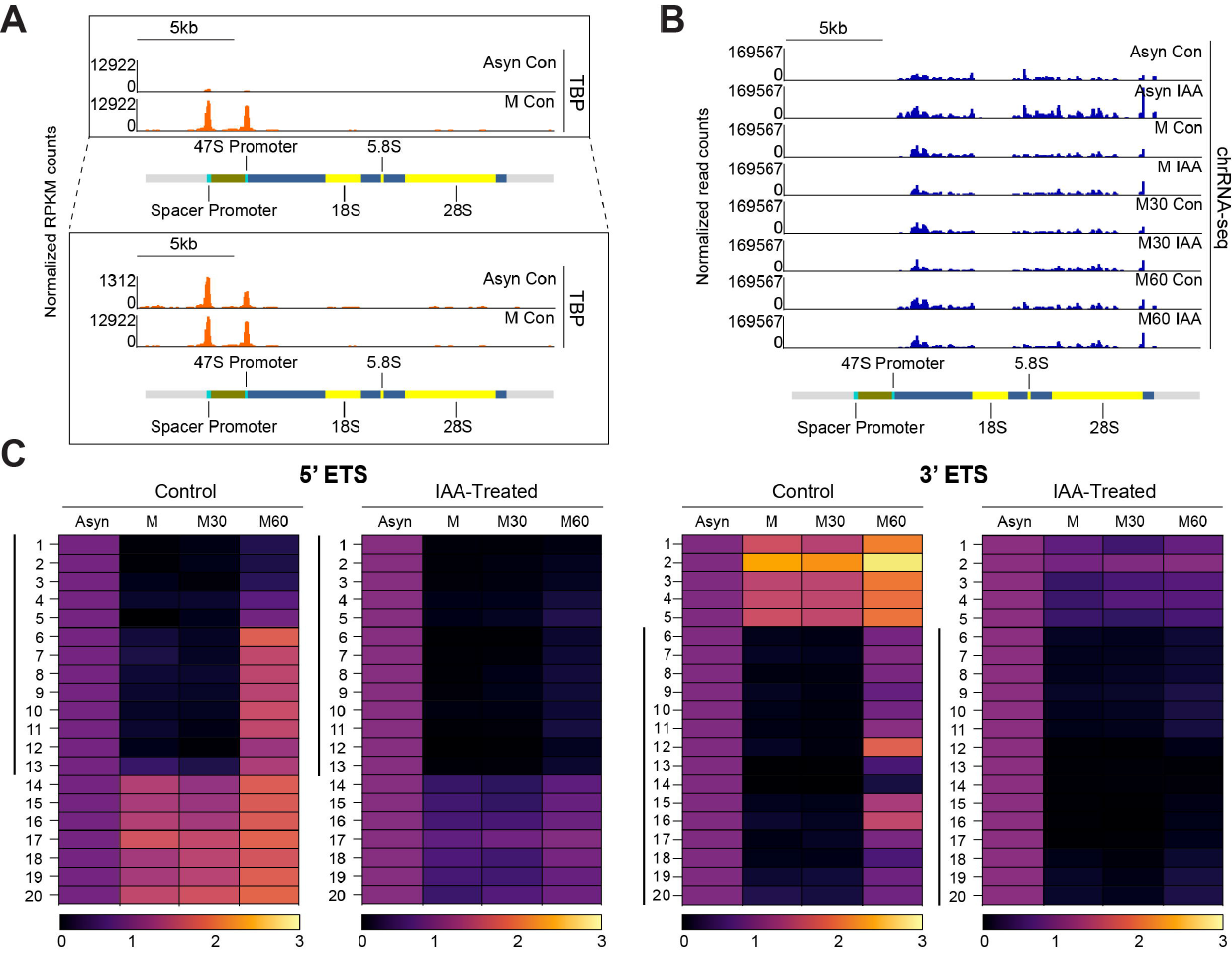
TBP bookmarks rDNA genes during Mitosis for efficient reactivation of rDNA transcription. **(A)** Gene browser tracks of the custom mm10-rDNA genome for TBP ChIP-seq in asynchronous control (Asyn Con) and nocodazole-treated (M Con) C64 mESCs for chR. Zoomed in view of the asynchronous track is shown below **(B)** Gene browser tracks of the custom mm10-rDNA genome for chrRNA-seq for control and IAA-treated mESCs during mitosis (M) 30 minutes after mitotic release (M30) and 60 minutes after mitotic release (M60). (**C**) Normalized heatmaps of chrRNA-seq for the custom mm10-rDNA 5’ ETS and 3’ ETS loci for control and IAA-treated C64 mESCs in asynchronous cells, during mitosis (M), 30 minutes after mitotic release (M30) and 60 minutes after mitotic release (M60). Regions were scaled to the same length, reads were binned by 100bp for a total of 20 bins (y-axis 1-20) and normalized to the non-nocodazole treated conditions (Ctrl or IAA). Bin regions marked within the lines represent bins that are affected by TBP depletion following mitotic reactivation.

How does mitotic TBP binding affect reactivation of rDNA genes following mitosis? To address this question, we reexamined previously published chrRNA-seq data of asynchronous (Asyn), mitotic (M), and 30 (M30) and 60 (M60) mins after release from mitotic arrest in both control and IAA-treated conditions. After realignment to the rDNA genome and normalizing to the spike-in control, we observe that in control conditions, certain regions exhibit a strong decrease in levels of chrRNA during mitosis, remain low 30 mins after mitotic release, and increase back to asynchronous levels 60 mins after mitotic release (Figure 4B). To further quantify the signal, we displayed the chrRNA-seq data as a heatmap for each annotated rDNA loci and observed that the intergenic regions display the greatest decrease in chrRNA levels in mitosis, followed by reactivation 60 mins after mitotic release (Figure S4B). Focusing on the intergenic regions, we segmented each region into 20 bins and displayed the average chrRNA-seq data per bin as a heatmap (Figure 4C). We observe a complete loss of chrRNA signal in certain bins for the 5’ETS (bins 1-13) and 3’ ETS (6-20) during mitosis, which then regain levels in 60 mins after mitotic release, demonstrating efficient mitotic arrest and release (Figure 4C). However, other regions displayed maintained or increased chrRNA levels during mitosis (Figure 4C, bins 14-20 in 5’ETS and bins 1-5 in 3’ETS, Figure S4B-C). Given that Pol I has been shown to be inactivated during mitosis, the lack of decrease in chrRNA signal in these regions during mitosis is likely indicative of contamination from highly abundant rRNAs.

We then examined the IAA-treated conditions in asynchronous, mitotic and 30 and 60 min after release from mitotic arrest. We observe a similar trend to control conditions where chrRNA levels decrease during mitosis and remain low 30 mins after mitotic release (Figure 4B). However, in the M60 IAA samples, we observe a distinct lack in transcription reactivation in certain regions compared to M60 control samples (Figure 4B), specifically in the intergenic regions (Figure S4B-C). The lack of reactivation following TBP depletion is more apparent in the 20-bin heatmaps of intergenic regions. The bins that experience a near complete loss of signal for the 5’ETS and 3’ETS during mitosis in control samples (bins 1-13 and 6-20 respectively) experience a delay in reactivation 60 mins after mitotic release compared to control, suggesting a specific role for TBP in Pol I transcription reactivation following mitosis (Figure 4C). Collectively, our results suggest that although TBP is not required for Pol I transcription of rRNAs, its bookmarking function may facilitate efficient reactivation of Pol I transcription after cell division.

## Discussion

In this study, we induced acute depletion of TBP via the mAID system and examined the role of TBP in Pol I transcription in mESCs. We report that the TBP paralog TRF2 does not functionally replace TBP for Pol I transcription as it does not bind to rDNA genes. We also report that, though Pol I occupancy is somewhat decreased upon TBP deletion, nascent rRNA production largely remains unaffected. Surprisingly, we find that mitotic TBP binding promotes efficient Pol I reactivation after mitosis

Despite sharing only ∼40% amino acid conservation to the core domain of TBP, TRF2 is predicted to also fold into a similar saddle-shaped structure (AlphaFold P62340) that is characteristic of TBP (25, 32). Historically, TRF2 has been shown to govern its own exclusive subset of Pol II genes when expressed in tandem with TBP, and in certain species it can activate transcription for TATA-less Pol II promoters (12, 33). In mESCs, we have shown that TRF2 has the ability to bind onto mRNA promoters in tandem with TBP, regardless of the presence of a TATA-box (8). Our results showing that TRF2 does not bind onto the rDNA promoter suggest that, unlike the diversity in Pol II pre-initiation complexes that accommodates paralogs of distinct subunits, the Pol I machinery has maintained high conservation by foregoing the presence of TBP paralogs.

Combined with previous studies, we show that, in the mESC system, TBP is not required for Pol I or Pol II transcription, but that Pol III transcription of tRNA genes is highly dependent on the presence of TBP. Through sequence and structural analyses, Pol I is believed to be more divergent from Pol II and Pol III initiation mechanisms (36–38). It is therefore intriguing to speculate why Pol I would be more similar to Pol II in their TBP-independent transcription compared to Pol III. In the case of Pol II, we previously showed that TFIID, consisting of 13 TAF subunits, forms in the absence of TBP, suggesting that the TAFs have compensatory roles in mammalian transcription (8). In Pol I initiation, TBP is also in a complex with five Pol I-specific TAFs (19–21), whereas in Pol III initiation, TBP is in complex with only two other factors (39). Perhaps the increased number of TAFs in Pol I and II initiation complexes provide additional redundancies, allowing for looser requirements for TBP in these systems.

Our findings that TBP is not required for general Pol I transcription, but plays a role in efficient reactivation after mitosis, are highly reminiscent of recent studies on the role of TBP in Pol II transcription. The role of TBP during mitosis is intriguing. Previous studies have shown that TBP is somewhat unique among TFs for having a long residence time on DNA, on the order of minutes (10, 42, 43), whereas most other TFs bind on the order of seconds (44). Perhaps such stable binding allows for TBP to overcome a transcriptionally repressive mitotic environment as the cells exit mitosis, providing a stable platform to assemble the initiation machinery in the critical transition period. Once the environment has fully transitioned into a more permissive state during G1, the advantage that TBP provides may no longer be necessary. In this way, the role of mammalian TBP in Pol I transcription may be highly context-dependent.

## Materials and Methods

### Key Resources Table

| Reagent type (species) or resource | Designation | Source or reference | Identifiers | Additional information |
| --- | --- | --- | --- | --- |
| Cell line (*Mus musculus*, male) | JM8.N4 mouse ES cells | KOMP repository | RRID: CVCL_J962 | Parental cell line used for all genetic manipulations |
| Cell line (*Mus musculus*, male) | mAID-TBP KI C94 | Teves et al. 2018 (10) | C94 | Endogenous knock-in of the minimal auxin inducible degron to N-terminus of TBP in JM8.N4 cells |
| Cell line (*Mus musculus*, male) | Halo-Pol II C64 | Teves et al. 2018 (10) | C64 | Endogenous knock-in of the HaloTag to C-terminus of Rbp1, largest subunit of Pol II, in mAID-TBP C94 cells |
| Cell line (*Mus musculus*, male) | TRF2 KO B8 | Kwan, Nguyen et al. 2023 (8) | B8 | Endogenous knock-out of the *Tbpl1* gene from Halo-Pol II C64 cells – see Materials and Methods |
| Cell line (*Mus musculus*, male) | TRF2-HA B8HA | Kwan, Nguyen et al. 2023 (8) | B8HA | Overexpressed HA-tagged TRF2 in B8 cells – see Materials and Methods |
| Antibody | α-TBP (Mouse, monoclonal) | Abcam | cat:ab51841 | Western blot (1:3000) |
| Antibody | α-Tubulin (Rabbit, polyclonal) | Abcam | cat:ab6046 | Western blot (1:7000) |
| Antibody | α-RPA1 (Mouse, monoclonal) | Santa Cruz | cat:sc-48385 | Western blot (1:2000)<br>CUT&Tag (1 μg) |
| Antibody | IRDye 800CW $\alpha$ -Mouse IgG (Goat polyclonal) | Licor | cat:926–32210 | Western blot (1:20000) |
| Antibody | IRDye 680RD α-Rabbit IgG (Goat polyclonal) | Licor | cat:926–68701 | Western blot (1:20000) |
| Peptide, recombinant protein | pAG-Tn5 | Epicypher | cat:15–1017 | Diluted to 1 x in each reaction |
| Data set | Pol I CUT&Tag | This study | GSE237001 |  |
| Data set | TBP CUT&Tag | Kwan, Nguyen et al. 2023 (8) | GSE206694 |  |
| Data set | TRF2 CUT&Tag | Kwan, Nguyen et al. 2023 (8) | GSE206694 |  |
| Data set | NET-seq | Kwan, Nguyen et al. 2023 (8) | GSE206695 |  |
| Data set | TBP ChIP-seq | Teves et al. 2018 (10) | GSE109962 |  |
| Data set | chrRNA-seq | Teves et al. 2018 (10) | GSE109963 |  |

### Cell Culture

For all experiments, the mouse ES cell line C64, B8, or B8HA was used. Cell lines were generated as previously stated (8, 10). ES cells were cultured on 0.1% gelatin-coated plates in ESC media KnockOut DMEM (Corning) with 15% FBS (HyClone), 0.1 mM MEM non-essential amino acids (Gibco), 2 mM GlutaMAX (Gibco), 0.1 mM 2-mercaptoethanol (Sigma-Aldrich) and 1000 units/ml of ESGRO (Chem-icon). ES cells were fed daily, cultured at 37 °C in a 5% CO_2_ incubator, and passaged every two days by trypsinization. For endogenously-tagged mAID-TBP C64 and B8 cells, TBP degradation was performed by addition of indole-3-acetic acid (IAA) at 500 μM final concentration to a confluent plate of cells for 6 hours.

### Western Blot Antibodies

Primary antibodies: α-TBP 1:3000 (Abcam ab51841), α-Tubulin 1:7000 (Abcam ab6046), and α-RPA1 (sc-48385). Secondary antibodies: IRDye 800CW Goat anti-mouse (Licor 926-32210), IRDye 800CW Goat anti-rabbit (Licor 925-32211), IRDye 680RD Goat anti-mouse (Licor 926-68070), or IRDye 680RD Goat anti-rabbit (Cedarlane 926-68701).

### Processing and alignment of NET-seq reads

NET-seq data was processed as previously described (8), but with the following modifications. Reads were trimmed and aligned to the custom mm10-rDNA genome (30) using STAR (v.2.7.3a). Read counts across the rRNAs were generated from normalized bam files using bedtools and analyzed with GraphPad Prism (45). The list of annotated Pol I rRNAs was obtained from the UCSC genome table browser with the following settings: clade: mammal, genome: mouse, assembly: mm10, group: all tables, database: mm10, table: rmsk, filter: repClass does match rRNA.

### Processing and alignment of chrRNA-seq reads

ChrRNA-seq data was processed as previously described (10), but with the following modifications. Reads were trimmed and aligned to the custom mm10-rDNA genome (30) using Tophat2. Bigwig files for IGV gene tracks were spike-in normalized and generated using DeepTools (46). Read counts across the rRNAs were generated from normalized bam files using bedtools and analyzed with GraphPad Prism (45). Heatmaps were generated using the DeepTools suite by scaling the regions and dividing the bins by 100bp.

### Processing and alignment of ChIP-seq reads

TBP ChIP-seq data was processed as described (10), but with the following modifications. Reads were aligned to the custom mm10-rDNA genome (30) using Bowtie2. Bigwig files for IGV gene tracks were RPKM normalized and generated using DeepTools (46). Replicates were merged using BigWigMerge.

### CUT&Tag Protocol

CUT&Tag was done as previously described (8), using the RNA Pol I antibody (sc-48385).

### Processing and alignment of CUT&Tag reads

TBP, TRF2-HA and Pol I CUT&Tag data was processed as described in Kwan and Nguyen 2023 but with the following modifications. Reads were aligned to the custom mm10-rDNA genome (30) using bowtie2 alignment. Bigwig files for IGV tracks were RPKM normalized and generated using DeepTools (46). Replicates were merged using BigWigMerge. Heatmaps were generated using the DeepTools suite. RPKM read count matrices were generated using DeepTools suite where RPKM normalized bigwig files were sorted into 10bp bins and rDNA loci (promoters and genes) were stretched to the same size using the scale-region command. RPKM counts were then averaged for each replicate and analyzed using GraphPad Prism.

## Acknowledgments

We thank R. Vander Werff and T. Stach (BRC-seq, UBC) for Illumina sequencing and S. Flibotte (LSI Bioinformatics facility, UBC) for implementation of NET-seq analyses. This work was supported by Life Sciences Institute Cores (LSI Imaging and ubcFLOW Core), supported by the UBC GREx Biological Resilience Initiative. S.S.T is a Canada Research Chair Tier 2 in Mechanisms of Gene Regulation, and is supported by the Michael Smith Foundation for Health Research.

## Author contribution

Conceptualization: JZJK, TFN, SST

Methodology: JZJK, TFN

Investigation: JZJK, TFN

Visualization: JZJK, TFN

Funding acquisition: JZJK, SST

Project administration:

Supervision: SST

Writing – original draft: JZJK, TFN, SST

Writing – review and editing: JZJK, TFN, SST

## Data and materials availability

All sequencing data have been deposited in Gene Expression Omnibus. All other data are available in the manuscript or in the supplementary materials. GEO IDs are in the Key Resources Table.

---

**Supplemental Figure 1.**
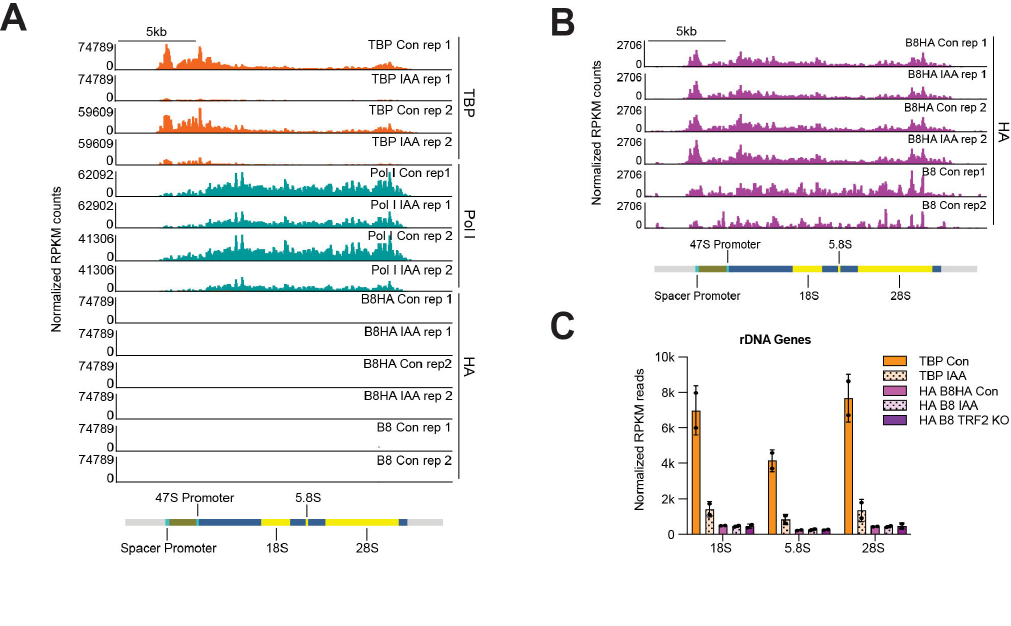
**(A-B)** Gene browser tracks of the custom mm10-rDNA genome chrR for TBP (orange), Pol I (teal) and TRF2-HA (magenta) in two biological replicates of control and IAA-treated C64 mouse embryonic stem cells (mESCs). Zoomed in gene browser track for TRF2-HA is shown in (B). **(C)** Average RPKM normalized read counts binned by 10bp at rDNA genes (18S, 5.8S and 18S) for TBP control (orange), TBP IAA (spotted light orange), HA B8HA control (magenta), HA B8HA IAA (spotted light magenta) and the negative HA B8 control (violet). Error bars represent standard deviation of 2 biological replicates.

**Supplemental Figure 2.**
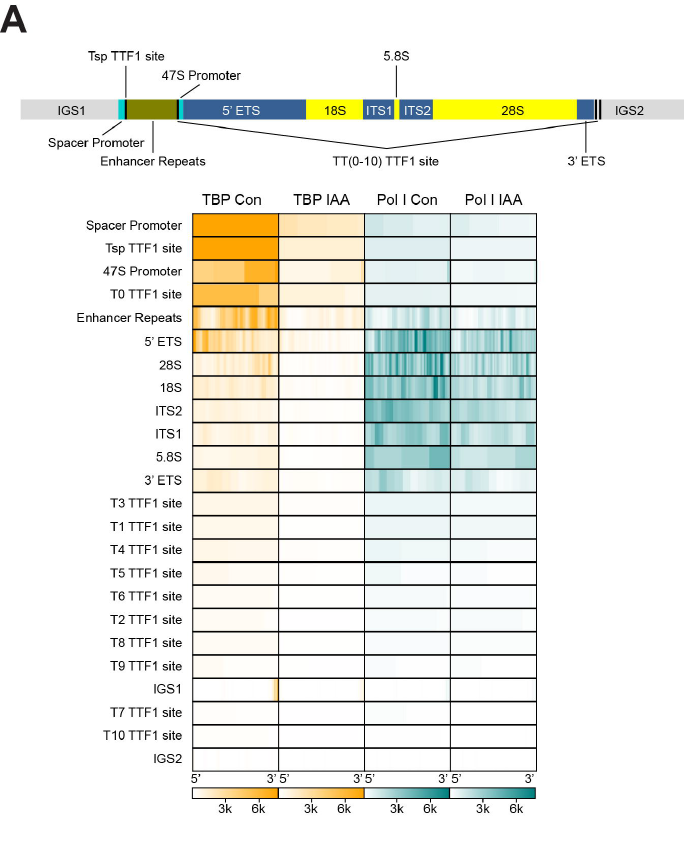
**(A)** RPKM normalized heatmaps of TBP CUT&Tag (orange) and Pol I CUT&Tag (teal) from the 5’ start to the 3’ end of the annotated custom mm10-rDNA regions binned by 10bp for control and IAA-treated C64 mESCs. Annotated chrR regions used in the heatmap are shown above.

**Supplemental Figure 3.**
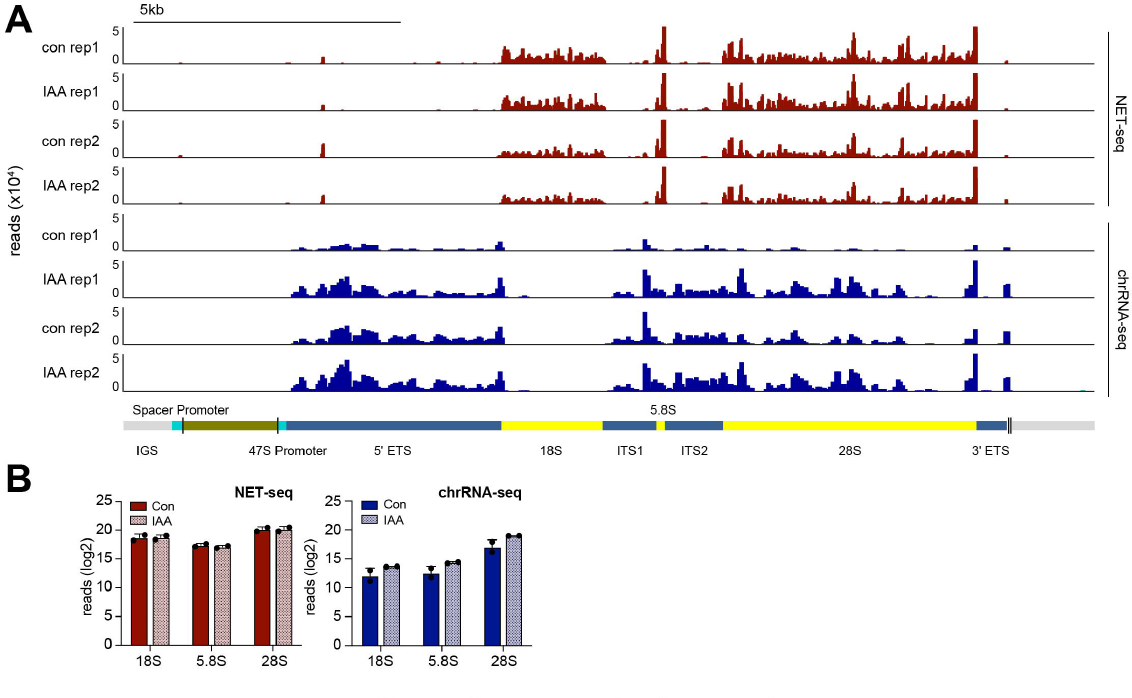
**(A)** Gene browser tracks of the custom mm10-rDNA genome for NET-seq (blue) and chrRNA-seq (cyan) in two biological replicates of control and indole-3-acetic acid (IAA)-treated C64 mouse embryonic stem cells (mESCs). **(B)** Normalized read counts of NET-seq (left) and chrRNA-seq (right) for control and IAA-treated mESCs across 18S, 5.8S, and 28S rRNAs. Error bars represent standard deviation of 2 biological replicates.

**Supplemental Figure 4.**
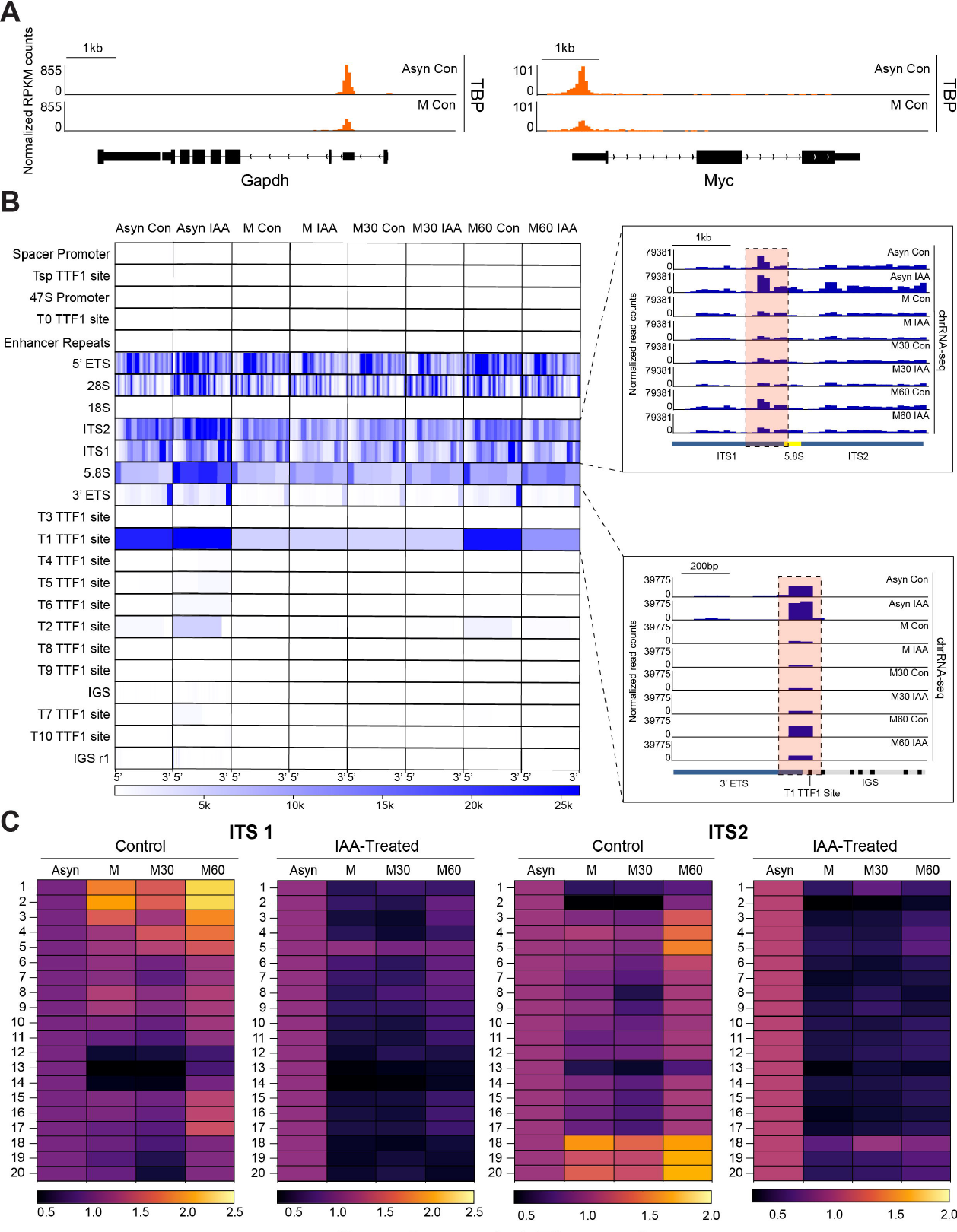
**(A)** Gene browser tracks of the custom mm10-rDNA genome for TBP ChIP-seq in asynchronous control (Asyn Con) and nocodazole-treated (M Con) C64 mESCs for *Gapdh* and *Myc*. **(B)** Normalized heatmap of chrRNA-seq from the 5’ start to the 3’ end of the of the annotated custom mm10-rDNA loci binned by 10bp for control and IAA-treated C64 mESCs during mitosis (M), 30 minutes after mitotic release (M30), and 60 minutes after mitotic release (M60). Boxes represent zoomed in regions of the indicated gene browser tracks for the custom mm10-rDNA genome. Highlighted regions in red show a failure to efficiently reactivate transcription 60 minutes after mitotic arrest. (M60 vs M60 IAA). **(C)** Normalized heatmaps of chrRNA-seq for the custom mm10-rDNA ITS1 and ITS2 loci for control and IAA-treated C64 mESCs during mitosis (M/MIAA), 30 minutes after mitotic release (M30/M30IAA) and 60 minutes after mitotic release (M60/M60IAA). Regions were scaled to the same length, reads were binned by 100bp for a total of 20 bins (y-axis 1-20) and normalized to the non-nocodazole treated conditions (Ctrl or IAA).

